# Allosteric modulation of GPCR-induced β-arrestin trafficking and signaling by a synthetic intrabody

**DOI:** 10.1101/2022.01.11.475811

**Authors:** Mithu Baidya, Madhu Chaturvedi, Hemlata Dwivedi-Agnihotri, Ashutosh Ranjan, Dominic Devost, Yoon Namkung, Tomasz Maciej Stepniewski, Shubhi Pandey, Minakshi Baruah, Bhanupriya Panigrahi, Jagannath Maharana, Ramanuj Banerjee, Jana Selent, Stephane Laporte, Terence E. Hebert, Arun K. Shukla

## Abstract

Agonist-induced phosphorylation of G protein-coupled receptors (GPCRs) is a primary determinant of β-arrestin (βarr) recruitment and trafficking. For several GPCRs, such as the vasopressin type II receptor (V_2_R), which exhibit high affinity for βarrs, agonist-stimulation first drives the translocation of βarrs to the plasma membrane, followed by endosomal trafficking. We previously found that mutation of a single phosphorylation site in V_2_R (i.e., V_2_R^T360A^) results in near-complete loss of βarr translocation to endosomes although βarrs are robustly recruited to the plasma membrane. Here, we show that a synthetic intrabody referred to as intrabody30 (Ib30), which selectively recognizes an active-like βarr1 conformation, rescues endosomal translocation of βarr1 for V_2_R^T360A^. In addition, Ib30 also rescues agonist-induced ERK1/2 MAP kinase activation for V_2_R^T360A^ to levels similar to that of the wild-type V_2_R. Molecular dynamics simulations reveal that Ib30 binding promotes active-like conformation in βarr1 with respect to the inter-domain rotation. Interestingly, we also observe that Ib30 enhances the interaction of βarr1 with β_2_-adaptin, which provides a mechanistic basis for the ability of Ib30 to promote endosomal trafficking of βarr1. Taken together, our data provide a novel mechanism to positively modulate the receptor-transducer-effector axis for GPCRs using intrabodies, which can potentially be integrated in the current paradigm of GPCR-targeted drug discovery.

**Significance:** The interaction of G protein-coupled receptors (GPCRs) with β-arrestins (βarrs) is a critical step in their regulatory and signaling paradigms. While intrabodies that bind to GPCRs, G proteins and βarrs have been utilized as biosensors and regulators of functional outcomes, allosteric targeting of receptor-transducer complexes to encode gain of function has not been documented so far. Here, we discover that a conformation-specific synthetic intrabody recognizing GPCR-bound βarr1 can allosterically enhance endosomal trafficking of βarr1 and agonist-induced ERK1/2 MAP kinase activation. This intrabody promotes an active-like βarr1 conformation and enhances the interaction of β_2_-adaptin with βarr1. Our findings establish a conceptual framework to allosterically modulate protein-protein interactions in GPCR signaling cascade to modulate their trafficking and signaling responses.

## Introduction

G protein-coupled receptors (GPCRs) recognize a broad spectrum of ligands and play a critical role in nearly every aspect of human physiology (1, 2). These receptors remain a major class of targets for novel drug discovery (3). The spatio-temporal aspects of GPCR signaling are tightly regulated by multifunctional β-arrestins (βarrs) (4, 5). Agonist-induced phosphorylation of GPCRs is a key determinant of βarr interaction and their ensuing functional outcomes (6, 7). While some GPCRs interact transiently with βarrs at the plasma membrane followed by rapid dissociation, others display a prolonged interaction resulting in endosomal trafficking of receptor-βarr complexes (8). These two patterns of βarr interaction and trafficking have been used to categorize the corresponding receptors as class A or class B GPCRs, respectively (8). Interestingly, distinct phosphorylation patterns on GPCRs have been linked to different βarr conformations, which in turn determine the resulting functional responses (9-11). While cumulative phosphorylation on GPCRs is typically believed to determine the affinity of βarr interaction, emerging evidence now suggests that spatial positioning of even single phosphorylation sites may make decisive contributions to βarr recruitment and subsequent functional outcomes (10, 11).

We previously reported that mutation of a single phosphorylation site in the vasopressin receptor (V_2_R) at Thr^360^ in the carboxyl-terminus (i.e., V_2_R^T360A^) dramatically altered βarr trafficking patterns (10). V_2_R is a class B receptor in terms of βarr interaction and trafficking where agonist-stimulation results in membrane recruitment of βarrs first followed by endosomal localization (8). Interestingly, upon agonist-stimulation of V_2_R^T360A^, βarrs efficiently translocate to the plasma membrane, but do not traffic to endosomal compartments unlike the wild-type receptor even after prolonged agonist-exposure (**Figure 1A**) (10). V_2_R^T360A^ also exhibits reduced levels of ERK1/2 activation compared to the wild-type receptor without any measurable effect on G protein-coupling as assessed by measuring cAMP production (10). This mutation leads to the disruption of a salt- bridge with Lys^294^ in βarr1 and consequently, reduces the fraction of active βarr1 conformation as assessed using molecular dynamics simulation (10).

**Figure 1.**
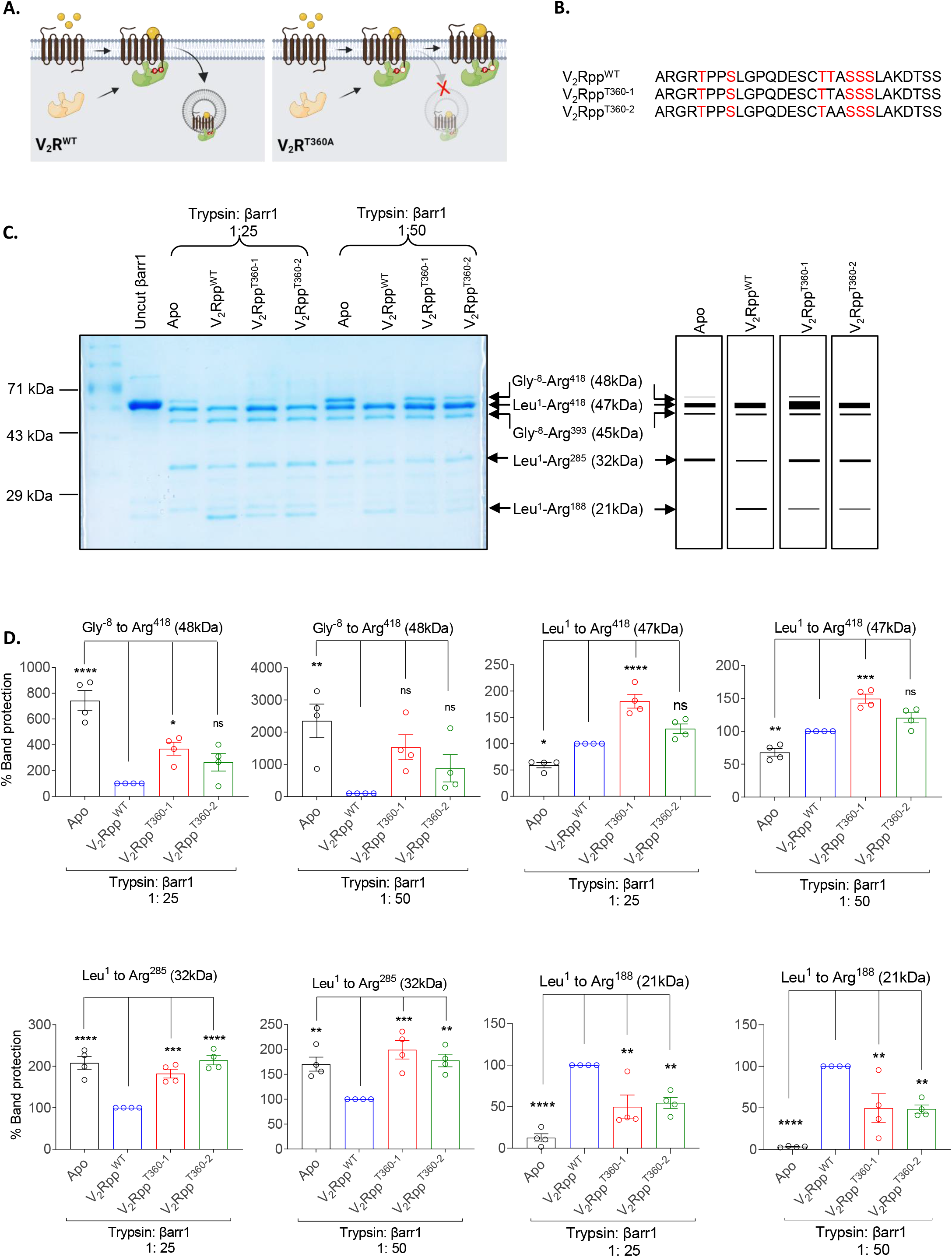
V_2_R^WT^ and V_2_R^T360A^ mutant impart distinct conformations to βarr1. **A** Schematic representation showing the inability of V_2_R^T360A^ to promote endosomal trafficking of βarr1. **B**. Sequences of the V_2_R^WT^ and Thr^360^ mutant phospho-peptides (V_2_Rpp^T360-1^ and V_2_Rpp^T360-2^) used. Phosphorylated S/T residues are highlighted in red. **C**. Limited trypsin proteolysis of V_2_Rpp^T360-1^/ V_2_Rpp^T360-2^ activated βarr1 show a band pattern distinct from both peptide-free βarr1 and V_2_Rpp^WT^ activated βarr1 indicating an intermediate conformation for the V_2_R^T360^ mutant. Free (Apo) or phosphopeptide-bound βarr1 was subjected to trypsin proteolysis at indicated trypsin: βarr1 ratio. The proteolysis reaction was quenched at 5min post digestion with SDS buffer and the digestion fragments were resolved on 12% SDS polyacrylamide gel. **D**. Densitometry based quantification of the % protection of individual tryptic fragments generated from βarr1 at indicated trypsin: βarr1 ratio is shown. Data from four independent experiments, normalized with respect to V_2_Rpp^WT^ condition, and analyzed using one-way ANOVA is presented here (*p<0.1, **p<0.01, ***p<0.001, **** p<0.0001).

In order to better understand this intriguing effect of Thr^360^Ala mutation in V_2_R, we set out to probe the conformation of βarr1 in complex with this receptor mutant, and compare it with the wild-type V_2_R, using a previously described intrabody30 (Ib30) based sensor (12). We observed that Ib30 robustly recognizes βarr1 recruited to the plasma membrane upon agonist-stimulation of V_2_R^T360A^. Surprisingly, we also found that Ib30 also promotes endosomal localization of βarr1 for V_2_R^T360A^ and rescues ERK1/2 MAP kinase activation to almost V_2_R^WT^ levels. We also discovered that Ib30 enriched the active-like conformational population of βarr1, and interestingly, also enhances the interaction of βarr1 with β_2_-adaptin. These findings establish the capability of Ib30 to allosterically modulate βarr1 trafficking and activation for V_2_R^T360A^, and potentially open a novel paradigm to modulate GPCR signaling using designer proteins.

## Results

### βarr1 conformations induced by V_2_Rpp^T360^ phospho-peptides

In order to probe whether the absence of Thr^360^ phosphorylation influences βarr1 conformation, we first synthesized two phospho-peptides corresponding to V_2_R^T360^ mutation, and used a previously described limited trypsin proteolysis assay (13) to compare βarr1 conformation induced by V_2_Rpp^T360^ phospho-peptides with that of V_2_Rpp^WT^. These two phospho-peptides, referred to as V_2_Rpp^T360-1^ and V_2_Rpp^T360-2^ contain a non-phosphorylated Thr or Ala at position 360, respectively, while the rest of the sequence and phosphorylation patterns are identical to V_2_Rpp (referred to as V_2_Rpp^WT^) (**Figure 1B**). Similar to a previous study (13), we observed that activation of βarr1 by V_2_Rpp^WT^ resulted in an accelerated cleavage of the 48kDa band (Gly^-8^-Arg^418^), protection of 47kDa and 45kDa bands (Leu^1^- Arg^418^ and Leu^1^-Arg^393^, respectively) and appearance of a 21kDa band (Leu^1^-Arg^188^) (**Figure 1C-D, Figure S1A-B**). Interestingly, we observed that V_2_Rpp^T360^ phospho-peptides also induced a proteolysis pattern qualitatively similar to that observed for V_2_Rpp^WT^. Still however, there were noticeable differences, including relatively slower proteolysis rates of the 47kDa band and weaker intensity of the 21kDa band. This observation indicates that V_2_R^T360^ phospho-peptides are capable of binding βarr1; however, they do not induce a fully-active βarr1 conformation as generated by V_2_R^WT^.

In order to further probe the conformation of βarr1 induced by V_2_Rpp^WT^ vs. V_2_Rpp^T360^ phospho-peptides, we measured the ability of conformationally-selective Fab30/ScFv30 sensors to recognize βarr1 conformation upon binding of these phospho-peptides by co-immunoprecipitation (co-IP) (**Figure 2A-B, Figure S2**). Fab30 and ScFv30 selectively recognize an active conformation of βarr1 induced by V_2_Rpp, and thus, have been used as conformational biosensors to monitor βarr activation *in vitro* (12, 14). We observed that Fab30/ScFv30 robustly interact with the V_2_Rpp^T360-1/2^- βarr1 complex albeit at lower levels than V_2_Rpp^WT^ (**Figure 2A-B, Figure S2**). We carried out this assay in presence of either 10-fold or 50-fold molar excess of the phospho-peptides compared to βarr1 but the reactivity patterns of Fab30/ScFv30 did not change significantly (**Figure 2A-B, Figure S2**). Similar to the limited proteolysis data presented in Figure 1, these data also suggest that V_2_Rpp^T360^ phospho- peptides induce a conformation in βarr1, which is qualitatively similar to that of V_2_Rpp^WT^ but not identical.

**Figure 2.**
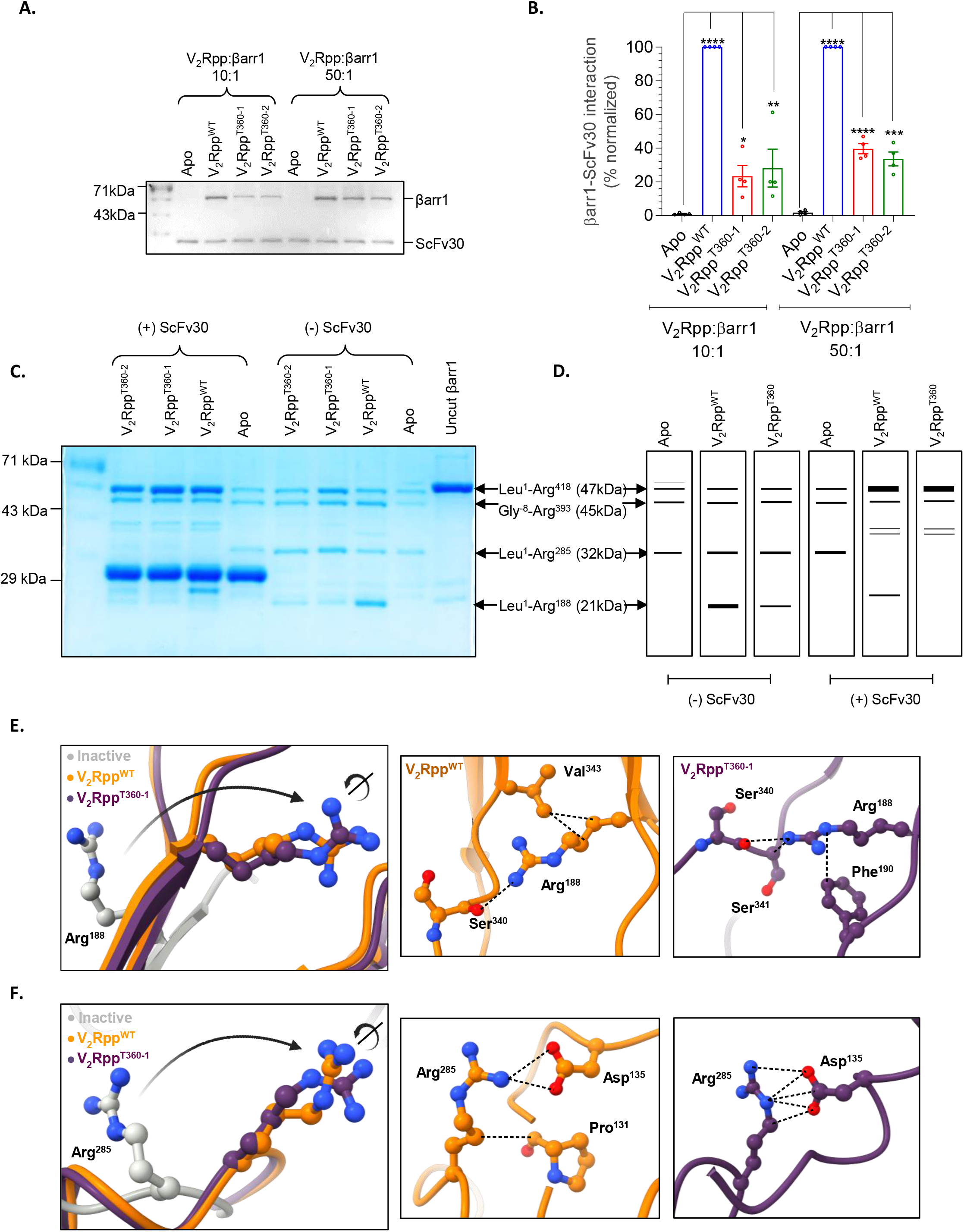
ScFv30 sensor efficiently recognizes βarr1 conformation upon stimulation of V_2_R^T360A^. **A**. Co-immunoprecipitation with ScFv30 shows that it recognizes the V_2_R^T360A^ bound βarr1. V_2_Rpp^T360-1^/ V_2_Rpp^T360-2^ activated βarr1 was incubated with ScFv30. The complex was subsequently pulled down with Protein-L agarose beads. A representative blot from four independent experiments is shown here. **B**. Densitometry-based quantification of βarr1-ScFv30 interaction normalized with V_2_Rpp^WT^- βarr1 control (taken as 100%) and analyzed using one-way ANOVA (*p<0.05, **p<0.01 ***p<0.001, **** p<0.0001) is shown. **C-D**. ScFv30 alters the tryptic digestion pattern of V_2_Rpp^T360-1^ or V_2_Rpp^T360-2^ bound βarr1. βarr1 activated with 50-fold molar excess of different phospho-peptides was subjected to limited trypsin proteolysis at a trypsin: βarr1 ratio of 1:50 in the presence or absence of ScFv30. The proteolysis reaction was quenched with SDS buffer after 30min and the digested fragments were separated by SDS-PAGE. The similarity in the digestion patterns of V_2_Rpp^WT^ and V_2_Rpp^T360-1^/ V_2_Rpp^T360A-2^ bound βarr1 in the presence of ScFv30 indicates that ScFv30 can indeed recognize the V_2_R^T360^ mutant activated βarr1. However, the differences in their digestion patterns also indicate that the V_2_R^WT^ bound conformation of βarr1 is not identical to that of the V_2_R^T360A^ bound βarr1. **E-F**. Structural snapshots comparing the relative orientation and local interaction networks of trypsin cleavage sites Arg^188^ and Arg^285^ in the crystal structures of basal (PDB: 1G4M, grey), V_2_Rpp^WT^ (PDB: 4JQI, orange) and V_2_Rpp^T360-1^ (PDB: 7DFA, violet) bound βarr1 is shown.

In order to further probe βarr1 conformation induced by different phospho-peptides, we carried out the limited proteolysis assay in presence of ScFv30 (**Figure 2C-D, Figure S3A**). We observed that the 47kDa band (Leu^1^-Arg^418^) is significantly protected in the presence of ScFv30, and the bands at 32kDa and 21kDa (Leu^1^-Arg^285^ and Leu^1^-Arg^188^, respectively) did not appear (**Figure 2C- D, Figure S3A**). Interestingly, the proteolysis patterns observed in presence of ScFv30 were nearly-identical for V_2_Rpp^WT^ and V_2_Rpp^T360^ phospho-peptides although an additional band at ∼30kDa was observed only with V_2_Rpp^WT^ (**Figure 2C-D**). Taken together with the co-immunoprecipitation data in **Figure 2A-B**, these data suggest that V_2_Rpp^T360^ phospho-peptides bind to βarr1 and ScFv30 potentially facilitates the transition of V_2_Rpp^T360^-bound βarr1 toward an active-like conformation.

In order to gain further insights into βarr1 conformation, we analyzed the crystal structures of βarr1 in basal, V_2_Rpp^WT^- and V_2_Rpp^T360^-bound states. As the distal carboxyl-terminus of βarr1 is not resolved in these structures, we focused primarily on Arg^285^ and Arg^188^, which are the trypsin cleavage sites yielding the 32kDa (Leu^1^-Arg^285^) and 21kDa (Leu^1^-Arg^188^) bands, respectively. Both of these residues exhibit a reorientation of their side chains between the apo and phospho-peptide-bound conformations (**Figure 2E-F**). Furthermore, an analysis of the local interaction networks of Arg^188^ and Arg^285^ calculated through CONTACT/ACT program within the CCP4 suite (15) revealed differences between the V_2_Rpp^WT^ vs. V_2_Rpp^T360^-bound conformations (**Figure 2E-F, Figure S3B**). These structural differences provide a plausible explanation for the proteolysis patterns obtained for V_2_Rpp^WT^ vs. V_2_Rpp^T360^, and support the hypothesis that V_2_R^T360^ induces an intermediate conformation in βarr1 compared to apo- and V_2_R^WT^-bound state.

### Intrabody30 rescues endosomal trafficking of βarr1 and ERK1/2 activation for V_2_R^T360A^

The experiments presented so far were carried out using isolated phospho-peptides *in vitro*. Thus, we set out to measure the reactivity of Ib30, an intrabody derived from Fab30, that efficiently recognizes active βarr1 in the cellular context, and also reports on βarr1 trafficking upon agonist-stimulation (11, 12). We first co-expressed SmBiT-βarr1 and LgBiT-Ib30 constructs with V_2_R^WT^ and V_2_R^T360A^ and measured agonist-induced changes in luminescence signal as a readout of βarr1-Ib30 interaction (**Figure 3A**). We observed a robust interaction of βarr1 and Ib30 upon agonist-stimulation of both, V_2_R^WT^ and V_2_R^T360A^, suggesting that the conformation of βarr1 is qualitatively similar in the cellular context, at least as reported by Ib30 sensor (**Figure 3B, Figure S4A**). We did not observe any measurable effect of Ib30 on G protein-mediated cAMP responses in case of V_2_R^T360A^, similar to V_2_R^WT^, suggesting that the intrabody does not significantly influence agonist-induced G_αs_ coupling (**Figure S5A-B**).

**Figure 3.**
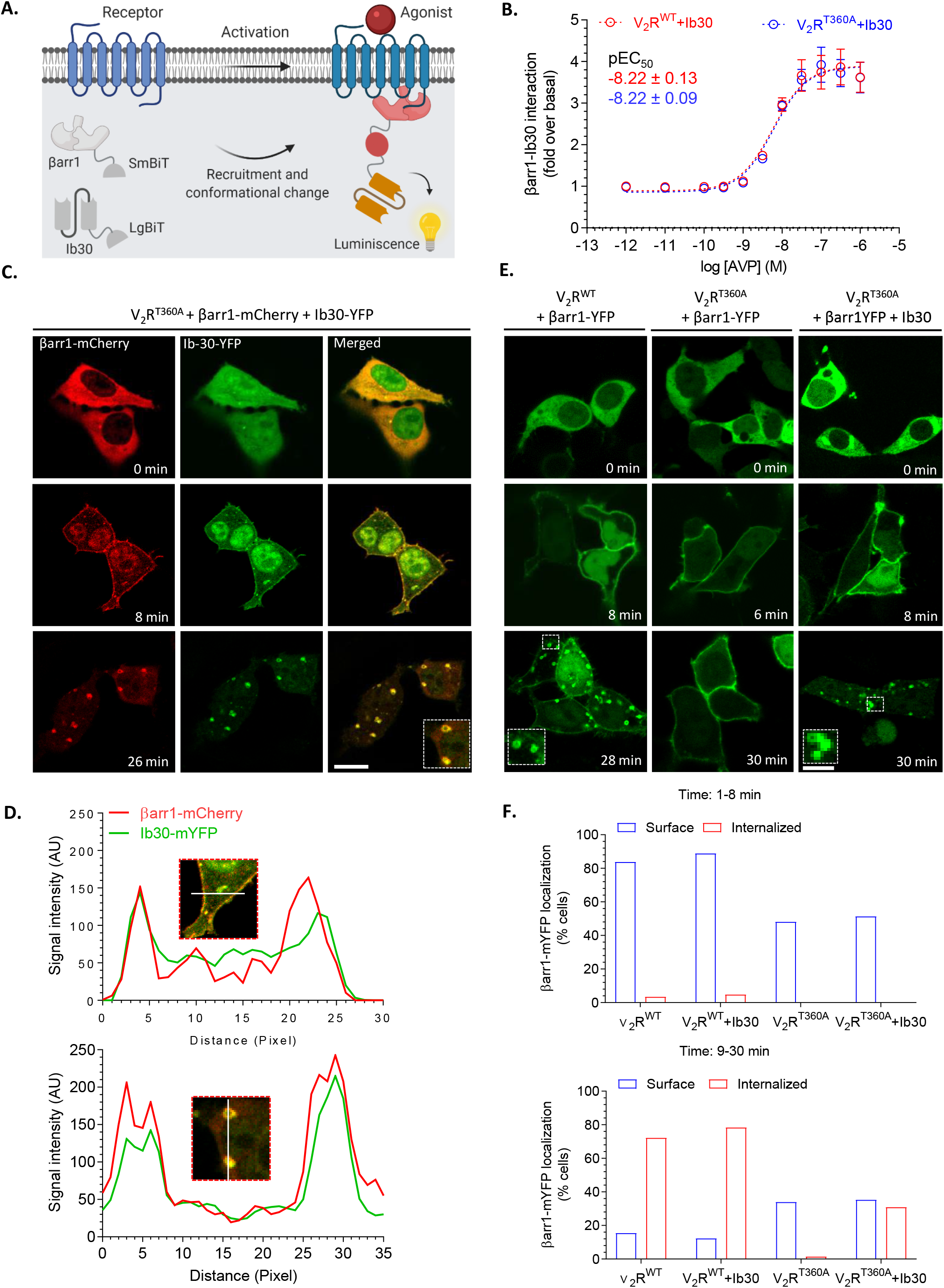
Intrabody30 (Ib30) sensor efficiently recognizes βarr1 conformation upon stimulation of V_2_R^T360A^. **A**. Schematic representation of NanoBiT-complementation-based Ib30 biosensor of βarr1 conformation, which selectively recognizes GPCR-bound βarr1. **B**. Ib30 exhibits efficient recognition of βarr1 upon stimulation of V_2_R^WT^ or V_2_R^T360A^. HEK-293 cells expressing the V_2_R^WT^ or V_2_R^T360A,^ together with SmBiT-tagged βarr1 and LgBiT-tagged Ib30 were stimulated with indicated doses of AVP followed by recording of luminescence signal. Data from three independent experiments are presented here. **C**. βarr1 and Ib30 are colocalized upon agonist-stimulation of V_2_R^T360A^. HEK-293 cells expressing V_2_R^T360A^ together with βarr1-mCherry and Ib30-YFP were stimulated with AVP (100nM) for indicated time-points followed by visualization using confocal microscopy. **D**. Line-scan analysis of the indicated regions from confocal micrographs confirms the colocalization of βarr1 and Ib30. **E**. Expression of Ib30 drives endosomal localization of βarr1 for V_2_R^T360A^. HEK-293 cells expressing V_2_R^WT^ or V_2_R^T360A^ together with βarr1-YFP were stimulated with AVP (100nM) and the localization of βarr1 was monitored using confocal microscopy. **F**. The effect of Ib30 on localization of βarr1 as assessed by manually scoring HEK-293 cells from multiple fields in three independent experiments. Captured confocal images were grouped in two classes i.e., 1-8min and 9-30min post-agonist stimulation to monitor membrane and endosomal localization, respectively. The bar graphs indicate the % of cells showing βarr localization at the surface or in endosomal punctate structures.

In order to directly visualize the ability of Ib30 to recognize βarr1 upon recruitment to V_2_R^T360A^, we co-expressed Ib30-mYFP construct together with βarr1-mCherry in HEK-293 cells expressing V_2_R^T360A^, and monitored the localization of βarr1 and Ib30 by confocal microscopy. Ib30 translocated to the plasma membrane upon agonist-stimulation, similar to βarr1, and exhibited robust colocalization with βarr1 (**Figure 3C-D**), in accord with the NanoBiT data presented in **Figure 3A-B**. Surprisingly however, we also observed that upon prolonged agonist-exposure (10-30min), both, βarr1 and Ib30 translocated to the endosomal vesicles and robustly co-localized (**Figure 3C-D**). This was unanticipated as βarr1 fails to translocate to endosomal vesicles even upon sustained agonist stimulation for V_2_R^T360A^ as reported previously (10). This observation led us to hypothesize that Ib30 may potentially modulate the trafficking pattern of βarr1 for V_2_R^T360A^ leading to endosomal localization of βarr1.

In order to test this, we co-expressed βarr1-mYFP construct in HEK-293 cells expressing either V_2_R^WT^ or V_2_R^T360A^ in presence or absence of HA-tagged Ib30. We monitored the localization of βarr1 in these cells upon agonist-simulation and scored the localization pattern of βarr1 in terms of plasma membrane vs. internalized vesicles. In line with data presented in **Figure 3C-D**, we observed that the presence of Ib30 indeed promotes endosomal trafficking of βarr1 for V_2_R^T360A^ (**Figure 3E-F**). βarr1 remains localized primarily at the plasma membrane even upon prolonged agonist-exposure in the absence of Ib30 as reported previously (**Figure 3E-F**) (10).

To further corroborate these findings, we used an intermolecular bystander BRET assay to monitor endosomal localization of βarr1 by using βarr1-R-Luc and GFP-FYVE constructs described previously (**Figure 4A**) (16). As shown in **Figure 4B**, we observed very low level of agonist-induced BRET for V_2_R^T360A^ in the presence of control intrabody (Ib-CTL) while V_2_R^WT^ exhibited a robust response as expected. Interestingly however, co-expression of Ib30 rescues the BRET signal (i.e., endosomal trafficking of βarr1) to almost the same level as V_2_R^WT^ (**Figure 4B**). We also observed an enhanced E_max_ in BRET assay for V_2_R^WT^ in the presence of Ib30, compared to Ib-CTL, although basal BRET was also slightly higher. Taken together with the confocal microscopy observations, these data establish that Ib30 not only recognizes V_2_R^T360A^-bound βarr1 conformation but also robustly promotes its trafficking to endosomal vesicles and thereby, rescuing the trafficking pattern of βarr1 similar to the wild-type receptor.

**Figure 4.**
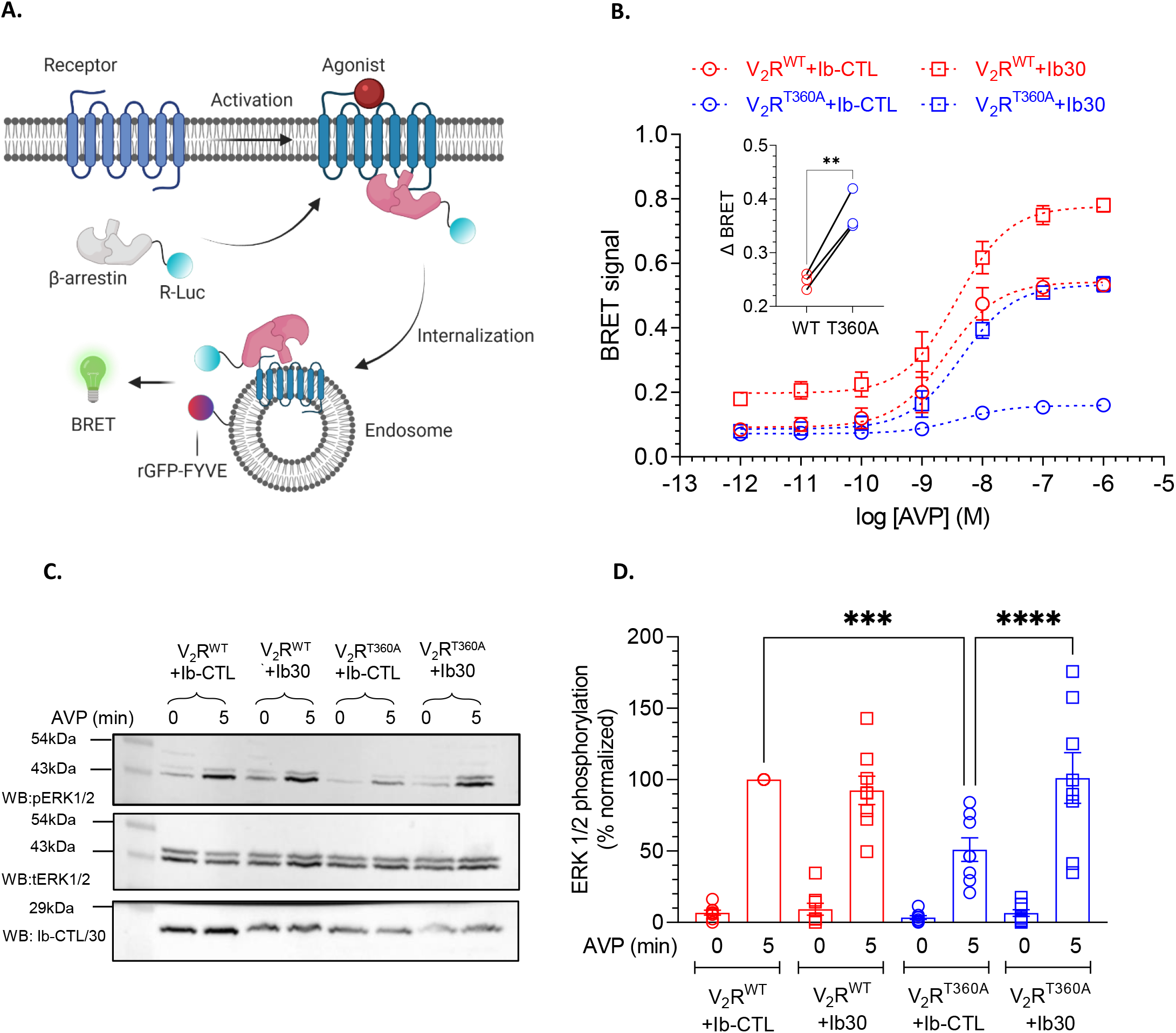
Effect of intrabody30 (Ib30) on endosomal trafficking and ERK1/2 activation for V_2_R^T360A^ mutant. **A**. Schematic representation of BRET-based endosomal localization assay for βarr1. **B**. Co- expression of Ib30 robustly promotes endosomal trafficking of βarr1 for V_2_R^T360A^ as assessed by BRET. HEK-293 cells expressing the V_2_R^WT^ or V_2_R^T360A,^ together with R-Luc-tagged βarr1 and Ib- CTL/Ib30 were stimulated with indicated doses of AVP followed by BRET measurement. *Inset* shows the change in BRET between the Ib-CTL and Ib30 conditions for V_2_R^WT^ and V_2_R^T360A^, respectively (**p<0.01, unpaired t-test). Data from three independent experiments are presented here. **C**. HEK- 293 cells expressing the indicated receptor construct together with Ib-CTL/Ib30 were stimulated with AVP (100nM) followed by detection of ERK1/2 phosphorylation using western blot. Expression of Ib-CTL/Ib30 is monitored using anti-HA antibody. **D**. Densitometry-based quantification from eight independent experiments, normalized with respect to V_2_R^WT^+Ib-CTL condition (treated as 100%), analyzed using one-way ANOVA is presented here (***p<0.001, **** p<0.0001).

We have previously reported that agonist-induced ERK1/2 MAP kinase activation is significantly attenuated for V_2_R^T360A^ compared to V_2_R^WT^. In order to measure whether Ib30 also modulates agonist-induced ERK1/2 activation, we compared ERK1/2 responses for V_2_R^WT^ and V_2_R^T360A^ with the control intrabody and Ib30 co-expression conditions. While Ib30 did not have a significant effect on ERK1/2 activation for the V_2_R^WT^, it robustly enhanced the level of phosphorylated ERK1/2 upon agonist-stimulation, nearly to that of the V_2_R^WT^ (**Figure 4C-D, Figure S4B**). Taken together with the endocytosis data, these findings demonstrate an allosteric effect of Ib30 to positively modulate βarr-mediated responses for the V_2_R^T360A^ mutant in the cellular context.

### Structural insights into βarr1 conformation and allosteric effect of Fab30

In order to better understand differences in the interaction pattern and binding mode of V_2_Rpp^WT^ vs. V_2_Rpp^T360^, if any, we analyzed the crystal structures of V_2_Rpp^WT^-βarr1 (PDB: 4JQI) and V_2_Rpp^T360-1^ (PDB: 7DFA). Interestingly, a segment of the V_2_Rpp^T360-1^ containing residues Pro^353^ to Thr^360^ shows a marked repositioning compared to the V_2_Rpp^WT^ binding pose (**Figure 5A**). In the V_2_Rpp^WT^-βarr1 crystal structure, pThr^360^ engages Lys^294^, Lys^11^ and Arg^25^ in βarr1 through ionic interactions, which is expectedly absent in case of V_2_Rpp^T360^ mutation. Of these, Lys^294^ in the lariat loop and Lys^11^ in the β- strand I of βarr1 are particularly noteworthy as they constitute a key part of the polar core and phosphate sensor, respectively. These interactions are critical in the process of βarr1 activation upon binding of phosphorylated carboxyl-terminus of GPCRs. Interestingly, pThr^359^ in V_2_Rpp^T360^ phospho- peptide engages with Lys^11^ but not with Lys^294^ or Arg^25^. This interesting structural rearrangement may in part explain an intermediate active-like conformation induced by V_2_Rpp^T360^ phospho-peptides as observed in limited proteolysis and ScFv30 co-IP assay.

**Figure 5.**
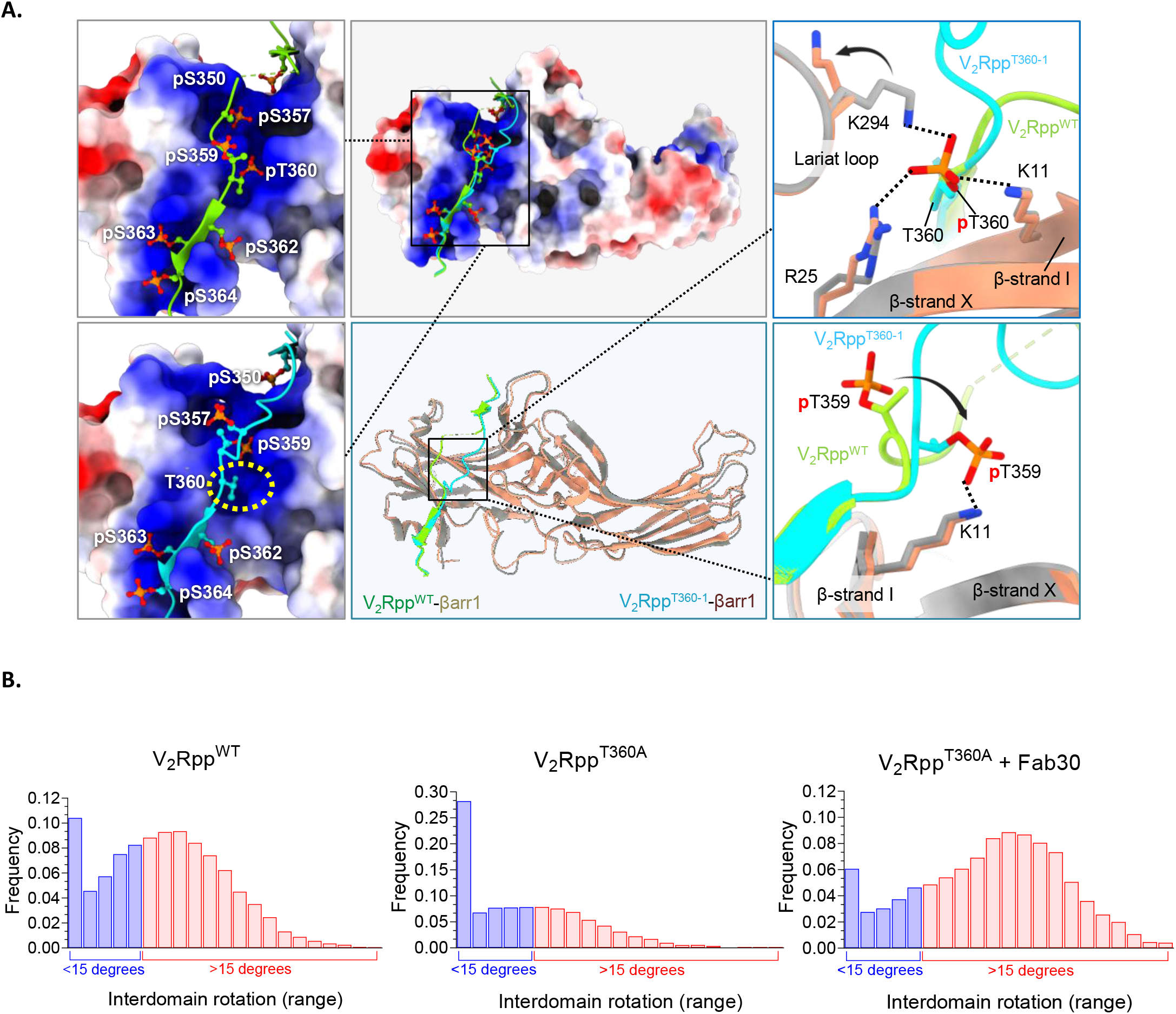
Intrabody30 (Ib30) stabilizes active conformation of βarr1. **A**. Structural snapshots of V_2_Rpp^WT^ (PDB: 4JQI) and V_2_Rpp^T360-1^ (PDB: 7DFA) bound βarr1 are shown. The superimposed structures display repositioning of the V_2_Rpp^T360-1^ N-terminal segment harbouring Thr^360^ residue (cyan) relative to the V_2_R^WT^ (green). Also, changes in ionic interactions of Thr^360^ with neighbouring residues are shown. For the V_2_R^WT^ bound βarr1, Thr^360^ engages with Lys^294^, Lys^11^ and Arg^25^. In the V_2_R^T360-1^ bound state, the Thr^360^ is non-phosphorylated and the side-chain of Thr^359^ is repositioned to interact with Lys^11^. **B**. MD simulation of βarr1 in complex with either V_2_Rpp^WT^ or V_2_Rpp^T360A^ based on crystal structure of V_2_Rpp-βarr1 (PDB: 4JQI) reveals an enrichment of inactive-like conformations of βarr1 in V_2_Rpp^T360A^-bound conformation. However, the binding of Fab30 to V_2_Rpp^T360A^-βarr1 complex robustly enriches the active-like conformational population of βarr1 as assessed by inter-domain rotation.

Next, in order to understand the effect of ScFv30 on βarr1 conformation in the context of V_2_R^T360^ mutation, we used molecular dynamics (MD) simulation on V_2_Rpp^WT^- and V_2_Rpp^T360A^-bound βarr1. We have previously reported that Thr^360^ to Ala mutation results in a significant shift in the population of βarr1 towards inactive-like conformation compared to the V_2_Rpp^WT^ as assessed in terms of the inter-domain rotation angle (10). In this study, we reproduce this behavior of V_2_Rpp^T360A^ compared to V_2_Rpp^WT^ demonstrating that introducing Thr^360^Ala mutation into the V_2_Rpp-βarr1 complex leads to a dramatic shift towards inactive-like conformations with an inter-domain rotation angles < 15° (**Figure 5B**, V_2_Rpp^WT^: 36% vs. V_2_Rpp^T360A^: 58%). Strikingly, simulation of the V^2^Rpp^T360A^- βarr1 complex in presence of Fab30 demonstrates that antibody binding increases the population of active-like βarr1 conformations (**Figure 5B**, V_2_Rpp^T360A^+Fab30: 80% vs. V_2_Rpp^T360A^: 42%). Taken together, these data underline the ability of ScFv30/Ib30 to promote active-like conformation in βarr1 in the context of Thr^360^Ala mutation, and provide a plausible mechanism for the positive allosteric effect of Ib30 on βarr1 trafficking to endosomes and ERK1/2 activation.

### Intrabody30 enhances βarr1-β_2_-adaptin interaction

Next, we set out to identify a potential functional correlate of Ib30-induced enrichment of active-like βarr1 conformation, and to reveal the mechanism of Ib30-mediated endosomal targeting of βarr1. As the interaction of βarrs with β_2_-adaptin is a prominent mechanism that drives GPCR endocytosis, we measured the effect of Ib30 on βarr1-β_2_-adaptin interaction. We used ear-domain of β_2_-adaptin (592-951) tagged with GST at the N-terminus and assessed its interaction with βarr1 in presence of V_2_R^T360A^. Here, we co-expressed βarr1 and V_2_R^T360A^ in HEK-293 cells, prepared cellular lysates after agonist stimulation, and used it for co-immunoprecipitation experiments. As presented in Figure 6A- B, we observed low but statistically significant interaction, compared to GST-alone control, between βarr1 and β_2_-adaptin in presence of Ib-CTL. Interestingly, the presence of Ib30 enhances this interaction several fold suggesting the ability of Ib30 to promote βarr1-β_2_-adaptin interaction (**Figure 6A-B**).

**Figure 6.**
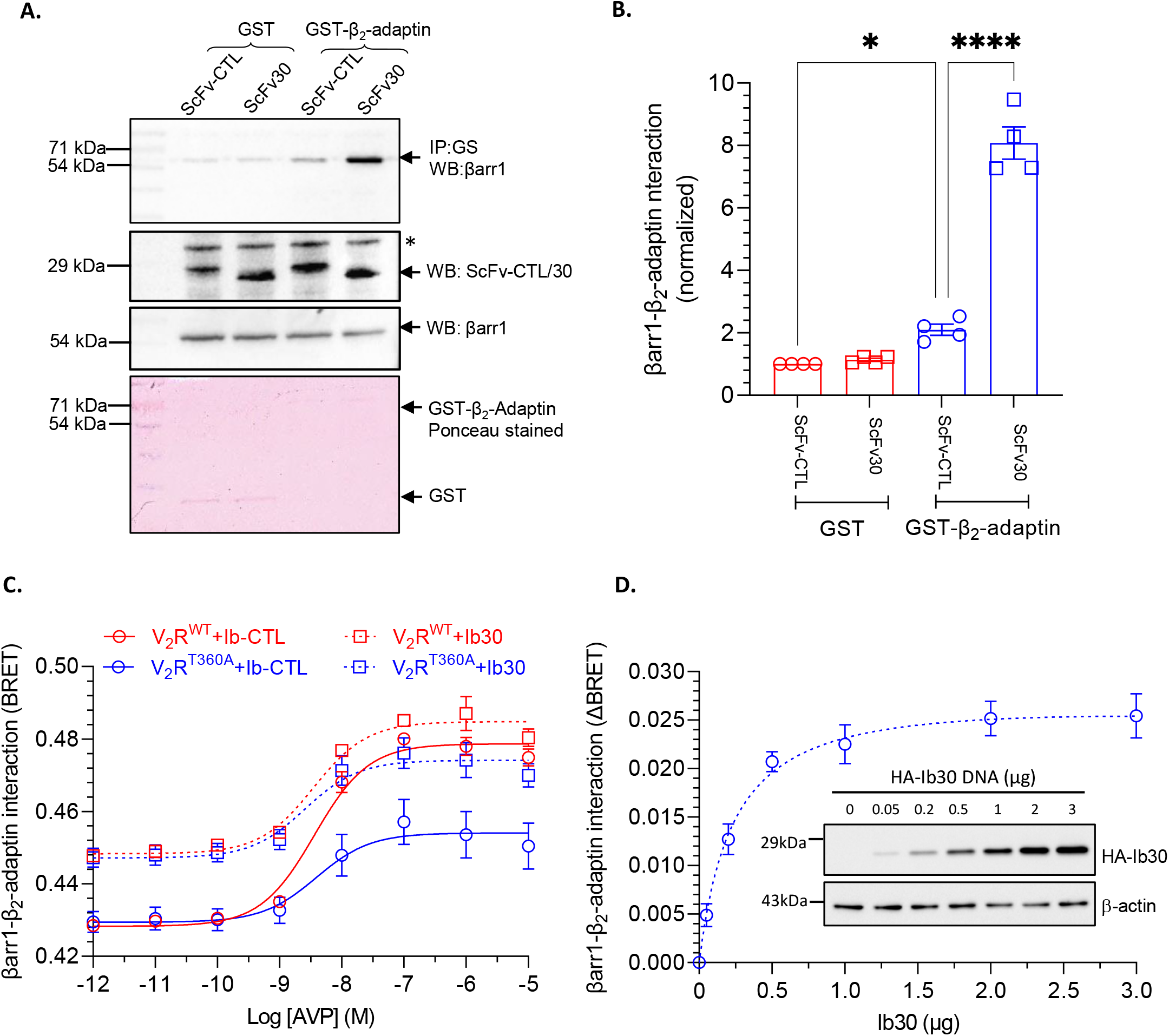
Intrabody30 (Ib30) enhances the interaction of β_2_-adaptin with V_2_R^T360A^-βarr1 complex. **A**. Purified GST-β_2_-adaptin (592-951) was incubated with V_2_R^T360A^ and βarr1 in presence of ScFv-CTL or SvFv30 followed by co-IP and Western blotting. Unconjugated GST was used as a negative control. A representative blot from three different experiments is shown here. The * symbol designates a non- specific band. **B**. Densitometry-based quantification of βarr1-β_2_-adaptin interaction from four independent experiments normalized with GST control and analyzed using one-way ANOVA (*p<0.05, ****p<0.0001). **C**. BRET between RlucII-tagged βarr1 and YFP-tagged β_2_-adaptin shows enhanced interaction between βarr1 and β_2_-adaptin in presence of Ib30, as compared to Ib-CTL, for both V_2_R^WT^ and V_2_R^T360A^. **D**. Ib30 induced increase in βarr1-β_2_-adaptin interaction exists even in the absence of either V_2_R^WT^ and V_2_R^T360A^. βarr1-β_2_-adaptin interaction in presence of Ib30 exhibits a concentration-dependent increase until saturating concentration of the latter. Data represent four independent experiments (mean±SEM). *Inset* shows a representative blot showing the concentration range of Ib30 used in the BRET experiment, normalized with respect to β-actin is shown.

In order to further corroborate this interesting finding in cellular context, we next used a previously described BRET-based assay (17) to monitor agonist-induced interaction of βarr1 with β_2_- adaptin in the absence and presence of Ib30 for V_2_R^WT^ and V_2_R^T360A^ (**Figure 6C**). There was robust interaction between βarr1 and β_2_-adaptin for the V_2_R^WT^ upon agonist-stimulation in the presence of control intrabody, while the response was significantly lower for V_2_R^T360A^, which is in line with significantly less endosomal trafficking of βarr1. Interestingly, co-expression of Ib30 significantly enhanced βarr1-β_2_-adaptin interaction for V_2_R^T360A^, bringing to almost to the same level as V_2_R^WT^, although basal BRET was also higher under Ib30 co-expression conditions for both, V_2_R^WT^ and V_2_R^T360A^. The enhanced BRET signal for V_2_R^T360A^ in presence of Ib30 provides a plausible mechanism for its ability to positively influence βarr1 trafficking to endosomes. Moreover, the elevated basal BRET in the presence of Ib30 is also intriguing and it may result from the propensity of Ib30 to enhance the interaction between βarr1 and β_2_-adaptin, even under basal conditions. In order to test this hypothesis, we carried out a titration experiment, where we expressed Ib30 at increasing levels and assessed βarr1-β_2_-adaptin interaction in the BRET assay. As presented in **Figure 6D**, increasing expression of Ib30 indeed enhanced BRET in a saturable manner suggesting the ability of Ib30 to promote basal interaction between βarr1 and β_2_-adaptin. Taken together, these observations provide a mechanistic basis of Ib30-induced allosteric modulation of βarr1 trafficking pattern observed for V_2_R^T360A^.

## Discussion

In this study, we demonstrate that a synthetic intrabody (Ib30) can allosterically modulate agonist-induced trafficking patterns of βarr1 for V_2_R^T360A^ lacking a key phosphorylation site in its carboxyl-terminus. Ib30 imparts this transition from a class A to a class B trafficking pattern by enriching the fraction of active-like conformational populations of βarr1, and allosterically enhancing βarr1-β_2_- adaptin interactions. Moreover, Ib30 also rescues the reduced ERK1/2 activation for V_2_R^T360A^ to levels induced by the wild-type receptor. A previous study has demonstrated a critical role of βarr-β_2_- adpatin interaction in βarr-mediated ERK1/2 activation for V_2_R using a small molecule inhibitor of this interaction (17). Therefore, an increase in βarr1-β_2_-adaptin interaction in presence of Ib30 may provide a plausible mechanism for its ability to rescue agonist-induced ERK1/2 activation for V_2_R^T360A^. However, additional mechanisms may also contribute to this intriguing observation, and it would be interesting to probe this further in subsequent studies. While previous studies have used intrabodies as biosensors of GPCR activation (18), βarr trafficking (12), inhibitors of GPCR endocytosis (19) and Gβγ signaling (20), the current study provides yet another application of intrabodies to modulate βarr trafficking and alter functional outcomes.

A recent study elegantly depicted the structures of βarr1 in complex with several different phospho-peptides derived from the carboxyl-terminus of V_2_R including that corresponding to V_2_R^T360- 1^ (21). Interestingly, the binding affinities of βarr1 to V_2_Rpp^WT^ and V_2_Rpp^T360^ are comparable although V_2_Rpp^T360^ exhibits a slightly altered binding mode compared to V_2_Rpp^WT^ in the crystal structures (21). Therefore, it is unlikely that distinct trafficking patterns of βarr1 for V_2_R^WT^ vs. V_2_R^T360A^ originate from an affinity difference, and therefore, potentially point towards a conformational mechanism underlying this phenomenon. This is in fact supported by MD simulation analysis of βarr1 structure in complex with V_2_Rpp^WT^ vs. V_2_Rpp^T360A^. Mutation of Thr^360^ to Ala results in a significant reduction of βarr1 conformational population in active-like state as assessed in terms of inter-domain rotation. Strikingly however, simulations in the presence of Fab30 showed a dramatic enrichment of active-like conformational ensembles.

In the current conceptual framework, GPCR-βarr interactions are typically conceived to be biphasic and involve the phosphorylated carboxyl-terminus of the receptor and the cytoplasmic face of the activated transmembrane bundle (6, 7, 22-25). Previous studies have visualized such partially-engaged and fully-engaged GPCR-βarr complexes and deciphered functional outcomes associated with these distinct conformations (23-25). A recent study using NMR spectroscopy demonstrated that Fab30 binding to partially-activated βarr1 facilitates additional conformational changes in βarr1 leading to a fully-activated conformation (26). Our study therefore draws an interesting parallel where Ib30 allosterically rescues a functionally-compromised βarr1 conformation to a functionally- competent conformation and corresponding cellular responses. It would be interesting to explore in future studies whether the effect of Ib30 observed here for V_2_R^T360A^ is somehow linked to the transition between the partially- and fully-engaged βarr conformations in complex with the receptor.

The paradigm of βarr-AP2 interaction through β_2_-adaptin in driving GPCR endocytosis through clathrin-mediated endocytosis is mostly conserved across GPCRs (5). Therefore, our study also raises the possibility of using Ib30 to modulate the βarr1 trafficking for other GPCRs, and decipher previously unknown functions. It is relevant to mention here that Ib30 efficiently recognizes βarr1 in complex with several native GPCRs, although it was selected from a phage display library using V_2_Rpp-βarr1 as the target (12, 22). An emerging paradigm suggests catalytic activation of βarrs where they may continue to generate functional outputs even after dissociation from activated receptors (27-29). It is therefore tempting to speculate if Ib30 may indeed recognize such conformational “memory” and may help its visualization in the cellular context as well as at high resolution using direct structural approaches. In case of wild-type V_2_R, agonist-stimulation promotes co-localization of the receptor, βarr1 and Ib30 in endosomal vesicles (12); however, this remains to be determined for the V_2_R^T360A^ mutant in presence of Ib30.

In summary, we demonstrate that agonist-induced trafficking of βarrs can be allosterically modulated using conformation-specific intrabodies targeting protein-protein interactions. These findings open a new paradigm for modulating GPCR signaling in the cellular context and discovering the interplay of distinct βarr functions.

## Acknowledgements

Research in A.K.S.’s laboratory is currently supported by the Senior Fellowship of the Wellcome Trust/DBT India Alliance (IA/S/20/1/504916) awarded to A.K.S., Department of Biotechnology (DBT) (BT/PR29041/BRB/10/1697/2018), Science and Engineering Research Board (EMR/2017/003804, SPR/2020/000408, and IPA/2020/000405), Council of Scientific and Industrial Research [37(1730)/19/EMR-II], Young Scientist Award from Lady Tata Memorial Trust, and IIT Kanpur. A.K.S. is an EMBO Young Investigator and Joy Gill Chair Professor. M.B. was supported by the National Post-Doctoral Fellowship of SERB (PDF/2016/002930) and Institute Post-Doctoral Fellowship of IIT Kanpur. H.D.-A. is supported by National Post-Doctoral Fellowship of SERB (PDF/2016/002893) and BioCare grant from DBT (BT/PR31791/BIC/101/1228/2019). M.C. is supported by a fellowship from CSIR [09/092(0976)/2017-EMR-I]. The work in T.E.H.’s laboratory was supported by a grant from the Canadian Institutes of Health Research (PJT-15698) and T.E.H. holds the Canadian Pacific Chair in Biotechnology. The work in S.A.L.’s laboratory was supported by the Canadian Institutes of Health Research: PJT-162368 and PJT-173504. J.S. is supported by the Instituto de Salud Carlos III FEDER (PI18/00094) and the ERA-NET NEURON & Ministry of Economy, Industry, and Competitiveness (AC18/00030). T.M.S. is supported by the National Science Centre of Poland, project number 2017/27/N/NZ2/0257. We thank Drs. Archana and Manish K. Yadav for help with co-IP experiments and protein purification, respectively.

## Authors’ contribution

MB carried out confocal microscopy, assisted in the NanoBiT assay, β_2_-adaptin interaction experiments using co-immunoprecipitation and ERK1/2 MAP kinase activation experiments; MC generated the receptor constructs, carried out the limited proteolysis experiment with AR, participated in β_2_-adaptin interaction experiments using co-IP, and performed ERK1/2 MAP kinase activation experiments with SP; HD-A carried out GloSensor assay with help from MC and NanoBit assay with MB, participated in β_2_-adaptin interaction experiments using co-IP; MBa and BP performed Fab30/ScFv30 co-IP assay; RB and JM carried out structural analysis of crystal structures; DD performed BRET experiments to monitor endosomal trafficking of βarr1 under the supervision of TEH; TMS performed MD simulation studies under the supervision of JS; YN performed βarr1-β_2_- adaptin BRET experiments under the supervision of SAL; all authors contributed in data interpretation and manuscript writing; TEH and SAL edited the manuscript; AKS coordinated and supervised the overall project.

## Conflict of interest

Authors declare that they have no conflicts of interest.

## Material and methods

### General reagents

Most chemicals and molecular biology reagents were purchased from Sigma-Aldrich unless mentioned otherwise. HEK-293 cells (ATCC; cat. no. CRL-3216) were maintained at 37°C under 5% CO_2_ in Dulbecco’s modified Eagle’s medium (Gibco; cat. no. 12800-017) supplemented with 10% FBS (Gibco; cat. no. 10270-106) and 100U ml^-1^ penicillin and 100μg ml^-1^ streptomycin (Gibco; cat. no. 15140-122). Cells were cultured in 10cm dishes (Corning; cat. no. 430167) at 37°C under 5% CO_2_ and passaged at 70 to 80% confluency using 0.05% trypsin-EDTA for detachment. *Sf9* cells (Expression Systems; cat. no. 94-001F) were maintained as suspension cultures in ESF 921 media (Expression Systems; cat. no. 96-001-01). Lauryl Maltose Neopentyl Glycol (LMNG) was purchased from Anatrace (cat. no. NG310).

### Construct design and expression plasmids

The expression constructs for the wild-type human V_2_R and V_2_R^T360A^ mutants have been described previously (10). Briefly, the cDNA coding for V_2_R^WT^ with an N-terminal HA signal sequence and FLAG tag was PCR amplified and cloned in a customized pcDNA 3.1 (+) vector. This construct was also cloned in pVL1393 vector for expression in *Sf9* cells. The Thr^360^ mutation was generated on the V_2_R^WT^ backbone using Q5 Site-Directed Mutagenesis Kit (NEB). The βarr1-mYFP plasmid used for confocal imaging experiments was obtained from Addgene (cat. no. 36916). βarr1-mCherry plasmid was a gift from Dr. Mark Scott, Institut Cochin, France. The plasmids encoding ScFv-CTL, ScFv30, Ib-CTL-HA, Ib30-HA and Ib30-YFP have been described previously (12, 19). The V_2_R^WT^ and V_2_R^T360A^ constructs were also fused with a 15 amino-acid flexible linker to the small subunit of NanoLuc i.e., SmBiT at its N-terminus. Similarly, Ib30 were N-terminally fused with LgBiT fragment in pCAGGS vector for NanoLuc complementation-based NanoBit assay. For *in-vitro* assays, i.e., trypsin proteolysis and ScFv30/Fab30 co-IP experiments, βarr1 was purified from BL21 cells by Glutathione Sepharose (GS) affinity chromatography. All the constructs were sequence verified (Macrogen). V_2_R agonist AVP (arginine-vasopressin) was synthesized by Genscript, and phospho-peptides V_2_Rpp^WT^, V_2_Rpp^T360-1^ and V_2_Rpp^T360-1^ were synthesized by the peptide synthesis facility at Tufts University. The construct for GST-tagged β_2_-adaptin (residues 592-951, Rat, isoform 2) in pGEX4T1 vector was received as a kind gift from Dr. Thomas Pucadyil (Pune, India).

### Limited trypsin proteolysis assay

To qualitatively assess the effect of different V_2_R phospho-peptides i.e., V_2_Rpp^WT^, V_2_Rpp^T360-1^ and V_2_Rpp^T360-2^ on βarr1 conformation, limited trypsin proteolysis of βarr1 in the presence or absence of these phospho-peptides was performed. The protocol for trypsin proteolysis of βarr1 has been described previously (13). Briefly, βarr1 (5-10μM) was incubated in the absence or presence of (50:1 molar ratio, phospho-peptide: βarr1) the phospho-peptides for 30min at 4°C. Thereafter, L-1- Tosylamido-2-phenylethyl chloromethyl ketone (TPCK) treated Trypsin (Sigma-Aldrich; cat. no. T1426) was added to the βarr1 phospho-peptide mixture at a ratio of 1:25 and 1:50 (w/w) and the samples were incubated at 37°C for 5min. In addition to the indicated ratio of trypsin: βarr1, other ratios like 1:10, 1:100 and 1:250 were also tried. At 1:10 ratio, βarr1 was completely digested while at lower trypsin concentrations the resolution of the digested fragments was poor. At each time point, 20μl of the reaction mix (5μg of βarr1) was withdrawn and transferred to a fresh microcentrifuge tube containing 5μl of 5x SDS loading buffer in order to quench the proteolysis reaction. The digested samples were separated on 12% SDS-polyacrylamide gels by electrophoresis to determine the effect of phospho-peptides on the digestion pattern of βarr1. Additionally, to study how ScFv30 affects the digestion pattern of βarr1 when activated with different phospho-peptides, a 50-fold molar excess of ScFv30 was added to the βarr1 samples prior to proteolysis. Samples without ScFv30 were used as reference for comparison. After proteolysis with a 1:50 ratio of trypsin: βarr1, the samples were quenched at 30min and resolved by SDS-PAGE as described earlier.

### Surface expression of receptor mutants

The surface expression of V_2_R^WT^ and V_2_R^T360A^ used in different cellular assays was measured by whole-cell surface ELISA. For this, HEK-293 cells transfected with either V_2_R^WT^ or V_2_R^T360A^ were seeded at a density of 0.2 million per well in a 24-well plate precoated with 0.01% poly-D-Lysine (Sigma-Aldrich; cat. no. P0899). After 24hr, cells were fixed with 4% (w/v) paraformaldehyde (pH 6.9) on ice for 20min and washed three times with 1× tris-buffered saline (TBS) buffer [150mM NaCl and 50mM Tris-HCl (pH 7.4)]. Subsequently, nonspecific sites were blocked with 1% bovine serum albumin (BSA; prepared in 1× TBS) for 90min, followed by the incubation of cells with horseradish peroxidase (HRP)-coupled anti-FLAG M2 antibody (dilution-1:5000; Sigma-Aldrich; cat. no. A8592), prepared in 1% BSA for 90min. Cells were then washed three times with 1% BSA in TBS, and 200μl of tetramethylbenzidine (TMB) ELISA substrate (Thermo Fisher Scientific; cat. no. 34028) was added to each well. Once blue color appeared in the wells, the reaction was stopped by transferring 100μl of the solution to a different 96-well plate already containing 100μl of 1M H_2_SO_4_. Absorbance was measured at 450nm in a multimode plate reader (Victor X4-Perkin-Elmer). For normalization of signal across different wells, cell density was estimated using Janus Green (Sigma-Aldrich; cat. no. 201677) staining. TMB solution was removed from the wells; cells were washed with 1×TBS followed by incubation with 0.2% (w/v) Janus Green for 20min. Thereafter, cells were washed three times with distilled water and 800μl of 0.5N HCl was added to each well. 200μl of this solution was used for measuring the absorbance at 595nm. Normalized surface expression of receptor constructs was calculated as the ratio of absorbance at 450nm and 595nm.

### Intrabody NanoBiT assay

Here, we measured the conformational change in ligand-induced βarr1 recognized by Ib30 using NanoBiT assay. Ib30 and βarr1 were N-terminally fused to LgBiT and SmBiT respectively with the 15- amino acid flexible linker and inserted into the pCAGGS plasmid. For NanoBiT assay, HEK-293 cells at a density of 3 million were transfected with V_2_R^WT^ or V_2_R^T360A^ receptor (5μg), LgBiT-Ib30 (5μg) and SmBiT βarr1 (2μg) using PEI (Polyethylenimine); 1 mg ml^-1^) as transfection agent at DNA: PEI ratio of 1:3. After 16-18hr of transfection, cells were harvested in PBS solution containing 0.5mM EDTA and centrifuged. Cells were resuspended in 3ml assay buffer (HBSS buffer with 0.01% BSA and 5mM HEPES, pH 7.4) containing 10μM coelenterazine (GoldBio; cat. no. CZ05) at final concentration. The cells were then seeded in a white, clear-bottom, 96-well plate at a density of 0.7-0.9 × 10^5^ cells per 100μl per well. The plate was kept at 37:C for 90min in the CO_2_ incubator followed by incubation at room temperature for 30min. Basal readings were taken in luminescence mode of a multi-plate reader (Victor X4-Perkin-Elmer). The cells were then stimulated with varying doses of ligand AVP ranging from 1pM to 1μM (6x stock, 20μl per well) prepared in drug buffer (HBSS buffer with 5mM HEPES, pH 7.4). Luminescence was recorded for 60min immediately after addition of ligand. The initial counts of 4-10 cycles were averaged and fold increase was calculated with respect to vehicle control (unstimulated values) and analyzed using nonlinear regression four-parameter sigmoidal concentration–response curve in GraphPad Prism software.

### Confocal microscopy

For visualizing the effect of intrabodies on βarr mediated receptor trafficking, HEK-293 cells were co-transfected with 3μg of either V_2_R^WT^ or V_2_R^T360A^ along with 2μg of βarr1-mYFP in presence or absence of 2μg of Ib30 with help of polyethylenimine (Polysciences; cat. no. 23966) reagent (21μl) in 10cm plates. Transfection was performed in FBS-deficient DMEM (Gibco; cat. no. 12800-017) after which cells were replaced with DMEM supplemented with FBS (Gibco; cat. no. 10270-106). Post 24hr, cells were seeded onto poly-D-lysine (Sigma-Aldrich; cat. no. P0899) precoated glass bottom confocal dishes (SPL Lifesciences; cat. no. 100350) at a density of 1 million per dish. Cells were allowed to adhere to confocal dishes for 24hr. The next day, cells were starved in FBS-deficient DMEM for 4hr and then stimulated with 100nM AVP and live cells were visualized under the confocal microscope (Zeiss LSM 710 NLO). The confocal microscope was equipped with a motorized XY stage along with a temperature and CO_2_ controlled platform. For visualizing Ib30 and βarr1 together, cells were transfected with βarr1-mCherry (2μg) and Ib30-mYFP (2μg) along with V_2_R^T360A^ (3μg). To excite mYFP, a multi-line argon laser source was used and for the mCherry, a diode pump solid state laser source was used. The emitted signal was detected with a 32x array GaAsP descanned detector (Zeiss). For related experiments all microscopic settings including laser intensity and pinhole slit were kept in the same range and for avoiding any spectral overlap between two channels filter excitation regions and bandwidths were adjusted accordingly. Images were acquired in line scan mode and were subsequently processed post imaging in ZEN lite (ZEISS) software suite. For quantifying βarr trafficking to either membrane or endosomes, confocal images were categorized into early (1 to 8min) and late time points (9 to 30min) post agonist stimulation. The cells with βarr1- mYFP fluorescence in the plasma membrane were scored as surface localized and the cells with punctate structures in the cytoplasm were scored as internalized. In cases where βarrs were seen in both, the membrane and in cytoplasmic punctate structures, cells having more than three puncta in the cytoplasm were scored under internalized category. Biological replicates were imaged at least three times independently on different days. Scored data from the cell count were plotted as percentage of βarr recruitment from more than 500 cells for each condition. To avoid any discrepancy in manual counting three different individuals counted the images in a blinded and cross-checked fashion. All data were plotted in GraphPad Prism software.

### Agonist-induced cAMP responses measured by GloSensor assay

To measure cAMP accumulation (as a readout for G protein activation), 50-60% confluent HEK-293 cells were co-transfected with either V_2_R^WT^ or V_2_R^T360A^ DNA (2μg), luciferase-based 22F cAMP biosensor construct (3.5μg) and Ib-CTL (2μg) or Ib30 (1μg) DNA. After 18-20hr of transfection, cells were washed with 1xPBS and treated with trypsin-EDTA (0.05%). Detached cells were harvested and centrifuged at 1,000 rpm for 10min, and the cell pellet was resuspended in 0.5mg ml^-1^ luciferin (GoldBio; cat. no. LUCNA) solution prepared in 1X HBSS buffer (Gibco; cat. no. 14065) containing 20mM HEPES (pH 7.4). Cells were then seeded at a density of 0.1-0.125 million per 100μl in 96 well white plate. The same pool of cells was also seeded side by side for surface expression by whole cell surface ELISA. The cells seeded in 96-well plate were incubated for 1.5hr in 5% CO_2_ followed by additional 30min at room temperature. Subsequently, the basal luminescence was recorded for 5 cycles using a plate reader (Victor X4-Perkin-Elmer), followed by addition of indicated concentrations of agonist AVP and luminescence was recorded for 1hr (30 cycles). Data were corrected for baseline signal and percent normalized with respect to maximal agonist concentration of V_2_R^WT^+Ib-CTL.

### BRET assay for βarr1 trafficking

HEK-293T cells (ATCC) were grown in complete culture media (DMEM high glucose (Wisent; cat. no. 319-015-CL) supplemented with 10% FBS (Wisent; cat. no. 098150) and penicillin/streptomycin (Wisent; cat. no. 450-201-EL) in a tissue culture incubator set at 37°C providing 5% CO_2_. The day before transfection, cells were plated into well of a 6-well plate (Thermo scientific; cat. no. 140675) at 4,00,000 cells per well. The next day, media was changed for DMEM high glucose supplemented with only 2.5% FBS and cells were transfected using PEI (Polysciences; cat. no. 23966) as follow: 1μg total DNA composed of 10ng of V_2_R^WT^ or V_2_R^T360A^, 25ng of βarr1-RlucII, 100ng of rGFP-FYVE and either 300ng Ib-CTL or 50ng Ib30 and DNA amount was completed with pcDNA3.1(+) was mixed with 3μl of a 1mg ml^-1^ PEI solution and added drop-wise to cells. The plate was put back in the incubator till the next day. 24hr post-transfection, cells were detached and re-plated at 60,000 cells per well into a poly-L-ornithine-coated (Sigma-Aldrich; cat. no. P3655) white 96-well plate (Thermo scientific; cat. no. 236105) in complete culture media and left to grow for another 24hr. Then, the 96-well plate was washed once with Kreb’s/HEPES solution (146mM NaCl, 4.2mM KCl, 0.5mM MgCl_2_, 1mM CaCl_2_, 5.9mM glucose, and 10mM HEPES buffer, pH 7.4) and 80μl of Kreb’s/HEPES was added per well. The plate was put back in the incubator for 2-3hr to allow cells to rest before BRET measurement. After the resting time, cells were stimulated for 15min at 37°C by adding 10μl of AVP (Sigma-Aldrich; cat. no. V9879) at different concentrations prepared in Kreb’s/HEPES. To assess BRET, 10μl of a 20μM coelenterazine 400A (GoldBio; cat. no. C-320) solution diluted in Krebs/HEPES was added 5min before the end of the stimulation period. BRET was then monitored by measuring 3 consecutive luminescence readings at both 410nm and 515nm using a Tristar2 plate reader (Berthold. Technologies GmbH & Co. KG). BRET was calculated as the emission at 515nm/emission at 410nm and the 3 values were averaged. BRET data were plotted as dose-response curves using GraphPad Prism 6.

### Effect of Ib30 on agonist induced ERK1/2 phosphorylation

To assess the effect of Ib30 on βarr mediated signaling downstream to V_2_R^WT^ and V_2_R^T360A^ mutant, agonist induced ERK1/2 phosphorylation was measured. For this, 60-70% confluent HEK-293 cells were co-transfected with 0.25μg of indicated V_2_R constructs and 1μg of HA-tagged Ib30. A control intrabody (Ib-CTL) that does not recognize receptor bound βarr1 was also transfected in parallel at levels comparable to Ib30 (3μg) to achieve normalized expression levels of both the intrabodies. 24hr after transfection, cells were seeded into six-well plates at a density of 1 million cells per well. The next day, cells were serum-starved in DMEM for 6hr and were then stimulated with 100nM AVP (agonist for V_2_R) for indicated time points. After stimulation for selected time points, the media was aspirated and the cells were lysed in 100μl of 2× SDS protein loading buffer. Cellular lysates were heated at 95°C for 15min, followed by centrifugation at 15,000 rpm for 15min. 10μl of samples were loaded per well and separated by 12% SDS-polyacrylamide gel electrophoresis. Phosphorylated ERK1/2 signal was detected by Western blotting using anti–phospho-ERK1/2 antibody (dilution- 1:5000; CST; cat. no. 9101) followed by reprobing of the blots with anti–total-ERK1/2 antibody (dilution-1:5000; CST; cat. no. 9102). The expression of Intrabody was confirmed by probing with anti-HA antibody (dilution-1:5000; Santa-Cruz; cat. no. sc-805). Signal on the western blots was detected using the ChemiDoc imaging system (Bio-Rad), and densitometry-based quantification was carried out using Image Lab software (Bio-Rad).

### Molecular dynamics simulations

Data without Fab30 was adapted from a previous study (10). To generate V_2_Rpp^WT^-βarr1, V_2_Rpp^360A^- βarr1, and V_2_R^T360A^-βarr1-Fab30 complexes, we used previously determined crystal structure (22). Missing fragments in the βarr1 and V_2_Rpp structures were modelled using the loop modeller module available in the MOE package (www.chemcomp.com). In Fab30 we maintained residues 5 to 108 of the light chain and residues 1 to 123 of the heavy chain. The complexes were solvated (TIP3P water) and neutralized using a 0.15 concentration of NaCl ions. System parameters were obtained from the Charmm36M forcefield (30). Simulations were carried out using the ACEMD3 engine (31). Both systems underwent a 20ns equilibration in conditions of constant pressure (NPT ensemble, pressure maintained with Berendsen barostat, 1.01325 bar pressure), using a timestep of 2fs. During this stage restraints were applied to the backbone. This was followed with 3 × 2μs of simulation for each system in conditions of constant volume (NVT ensemble) using a timestep of 4fs. This allowed us to amass a total of 6μs simulation time per system. For each of the simulations we used a temperature of 310K, which was maintained using the Langevin thermostat, hydrogen bonds were restrained using the RATTLE algorithm. Non-bonded interactions were cut-off at a distance of 9Å, with a smooth switching function applied at 7.5Å. The inter-domain rotation angle of βarr1 was analysed using a script kindly provided by Naomi Latoracca (32).

### Co-immunoprecipitation (co-IP) assay

Co-IP was performed to evaluate the interaction between V_2_Rpp^WT^, V_2_RppT^360-1^ and V_2_Rpp^T360-2^ with βarr1 in presence of Fab30 and ScFv30. 5μg of purified βarr1 was activated with 10-fold and 50-fold molar excess of phospho-peptides for 1hr at room temperature (25 °C) in binding buffer (20mM HEPES, pH7.4, 100mM NaCl). Thereafter, the activated βarr1 was incubated with 2.5μg of purified Fab30 or ScFv30. Subsequently, 20μl of pre-equilibrated Protein L beads (GE Lifesciences; cat. no. 17547802) were added to the reaction mixture and incubated for an additional 1hr at room temperature, which was followed by extensive washing (3–5 times) with binding buffer + 0.01% LMNG. Elution was taken with 2X SDS loading buffer. Interaction of Fab30 and ScFv30 with βarr1 in presence of phospho-peptides was visualized using Coomassie staining of the gels. Band intensity was analysed by ImageJ gel analysis software.

To assess the effect of ScFv30 on V_2_R^T360A^ induced βarr1-β_2_-adaptin interaction, we performed co-immunoprecipitation assay (co-IP). The V_2_R^T360A^ receptor was expressed in *Sf9* cells, stimulated with 100nM AVP and centrifuged to obtain receptor pellet. The receptor pellet was resuspended in appropriate volume of lysis buffer having 20mM HEPES, 150mM NaCl, 1X PhosSTOP (Roche; cat. no. 04906837001) and 1X protease inhibitor (Roche; cat. no. 04693116001), subjected to Dounce homogenization and incubated with 1μg of purified βarr1 for 30min at room temperature. The receptor-βarr complex was again incubated with 5μg of purified ScFv30 or ScFv- CTL for another 30min and solubilized with 1% LMNG for 1hr. Meanwhile, GST or GST-β_2_-adaptin protein (2.5μg) was immobilized on 20μl buffer (20mM HEPES, 150mM NaCl) equilibrated GS beads (1hr at room temperature) and washed once to remove any unbound protein. Subsequently, the supernatant from solubilized complex was allowed to bind with protein bound GS beads (1hr at room temperature) followed by three washes with wash buffer (20mM HEPES, 150mM NaCl, 0.01% LMNG). The bead-bound complex was eluted in 2X SDS loading buffer. Eluted samples were separated by 12% SDS–polyacrylamide gel electrophoresis and probed using βarr antibody (dilution- 1:10000; CST; cat. no. 4674). After solubilization, 20μl of lysate was set aside for confirming equal loading of βarr1 and ScFv. The lysate was run on separate 12% SDS–polyacrylamide gel and probed using βarr antibody and HRP-coupled protein L antibody (dilution-1:2,000; GenScript; cat. No. M00098) by western blotting. Band intensity was analysed by Image Lab software (Bio-Rad).

### BRET assay for βarr1-β_2_-adaptin interaction

To monitor βarr1 and β_2_-adaptin interactions, BRET assays between βarr1-RlucII and β_2_-adaptin-YFP were performed as described (17). HEK-293 cells were seeded at a density of 1×10^6^ cells per 100mm dish and transfected the next day with 250ng of V_2_R^WT^ or V_2_R^T360A^ along with 120ng of βarr1-RlucII, 1μg of β_2_-adaptin-YFP, and either 1.5μg Ib-CTL or 1μg Ib30 using PEI. Briefly, a total 6μg of DNA (adjusted with pcDNA3.1/zeo(+)) in 0.5ml of PBS was mixed with 12μl of PEI (25kDa linear, 1mg ml^-1^) in 0.5ml PBS and then incubated for 20min prior to applying to the cells. After 24hr, cells were detached and seeded onto poly-ornithine-coated 96-well white plates at a density of ∼35,000 cells per well for the BRET assays, which were performed 48hr after transfection. For BRET assays, cells in 96-well plates were washed once with Tyrode’s buffer (140mM NaCl, 2.7mM KCl, 1mM CaCl_2_, 12mM NaHCO_3_, 5.6mM D-glucose, 0.5mM MgCl_2_, 0.37mM NaH_2_PO_4_, 25mM HEPES, pH 7.4) and left in Tyrode’s buffer for 1h at room temperature. Cells were stimulated with various concentrations of AVP for 45min then BRET signals were measured using a plate reader (Victor X4-Perkin-Elmer). Coelenterazine h (Nanolight™, final concentration of 5μM) was added 25min prior to BRET measurement. The filter set used was 460/80nm and 535/30nm for detecting the RlucII, *Renilla* luciferase (donor) and YFP (acceptor) light emissions, respectively. The BRET ratio was determined by calculating the ratio of light emitted by YFP over light emitted by RlucII. For Ib30 titration, HEK- 293 cells were transfected with 120ng of βarr1-RlucII and 1μg of β_2_-adaptin-YFP along with various amounts (0 to 3μg) of Ib30 in 100mm dishes or scale down to 1/6 in a well in 6 well plates. BRET signals were measured in absence of ligand stimulation. Expression levels of Ib30 were accessed by western blotting with anti-HA-peroxidase conjugate (dilution-1:1000, Sigma-Aldrich; cat. no. 12013819001). Anti-β-actin antibody (dilution-1:2000, Santa Cruz Biotechnology; cat. no. sc-47778) was used for a loading control.

### Data quantification and statistical analysis

The experiments were conducted at least three times and data (mean ± SEM) were plotted and analyzed using GraphPad Prism software (Prism 8.0). The data were normalized with respect to proper experimental controls and appropriate statistical analyses were performed as indicated in the corresponding figure legends.

## Notes

### Competing Interest Statement

The authors have declared no competing interest.

