## Supplemental figures 1-5 for "Allosteric modulation of GPCR-induced β-arrestin trafficking and signaling by a synthetic intrabody"

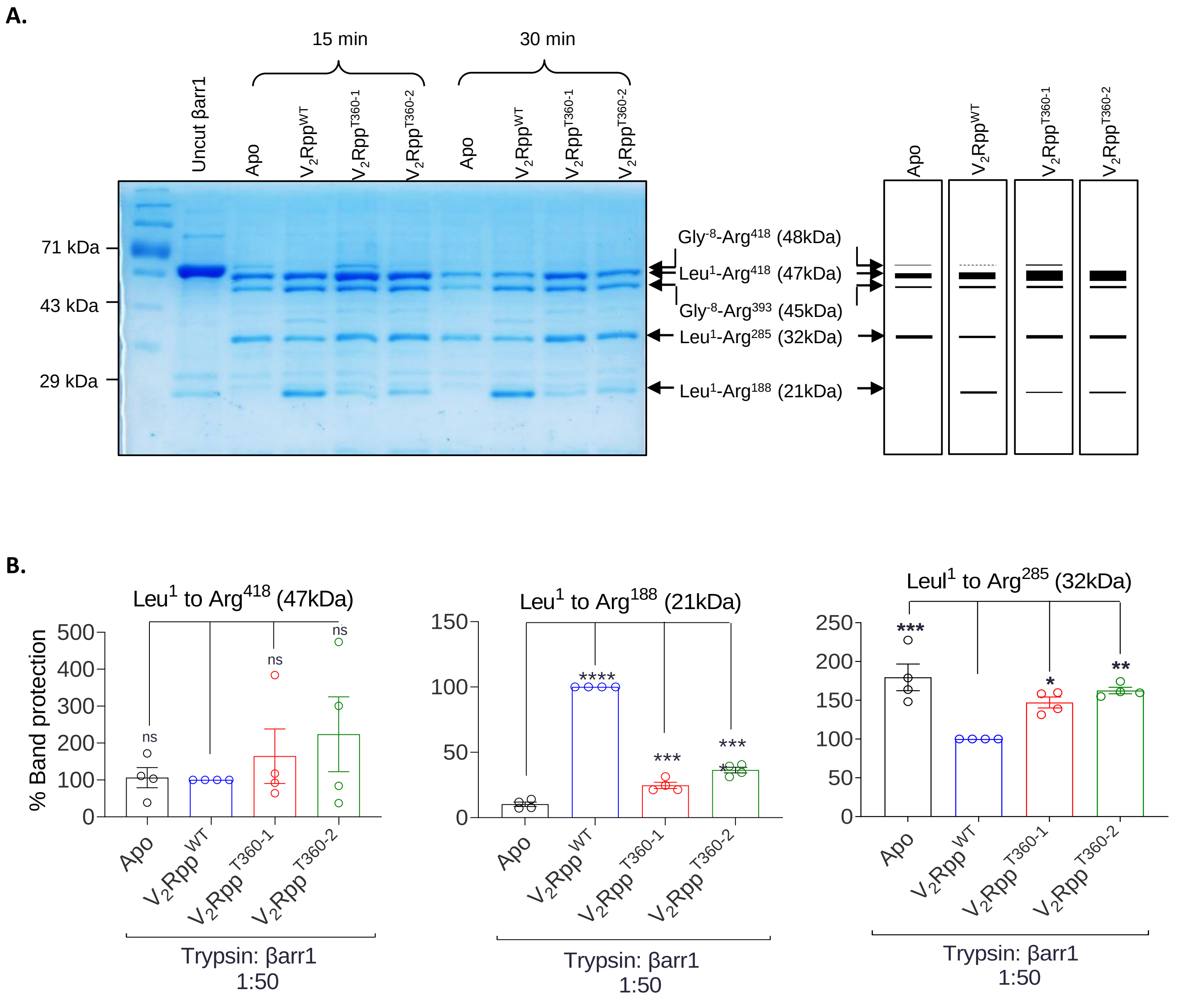


**Figure S1. Limited trypsin proteolysis of βarr1: A.** The trypsin digestion pattern of $V_2Rpp^{WT}$, $V_2Rpp^{T360\text{-}1}$ or $V_2Rpp^{T360\text{-}2}$ bound βarr1 is shown at indicated time points. βarr1 activated with 50-fold molar excess of different phosphopeptides was subjected to limited trypsin proteolysis at a trypsin: βarr1 ratio of 1:50. The proteolysis reaction was quenched with SDS buffer after 30min and the digested fragments were separated by SDS-PAGE. There are evident differences in the fragments ($Leu^1$-$Arg^{418}$ and $Leu^1$-$Arg^{188}$) resulting from digestion of ligand-free and ligand bound ($V_2Rpp^{WT}$, $V_2R^{T360\text{-}1}$ or $V_2Rpp^{T360\text{-}2}$) βarr1. **B.** Densitometry based quantification of the % protection of individual tryptic fragments generated from βarr1. Data from four independent experiments, normalized with respect to $V_2R^{WT}$ condition, and analyzed using one-way ANOVA is presented here (*p<0.1, **p<0.01, ***p<0.001, **** p<0.0001).

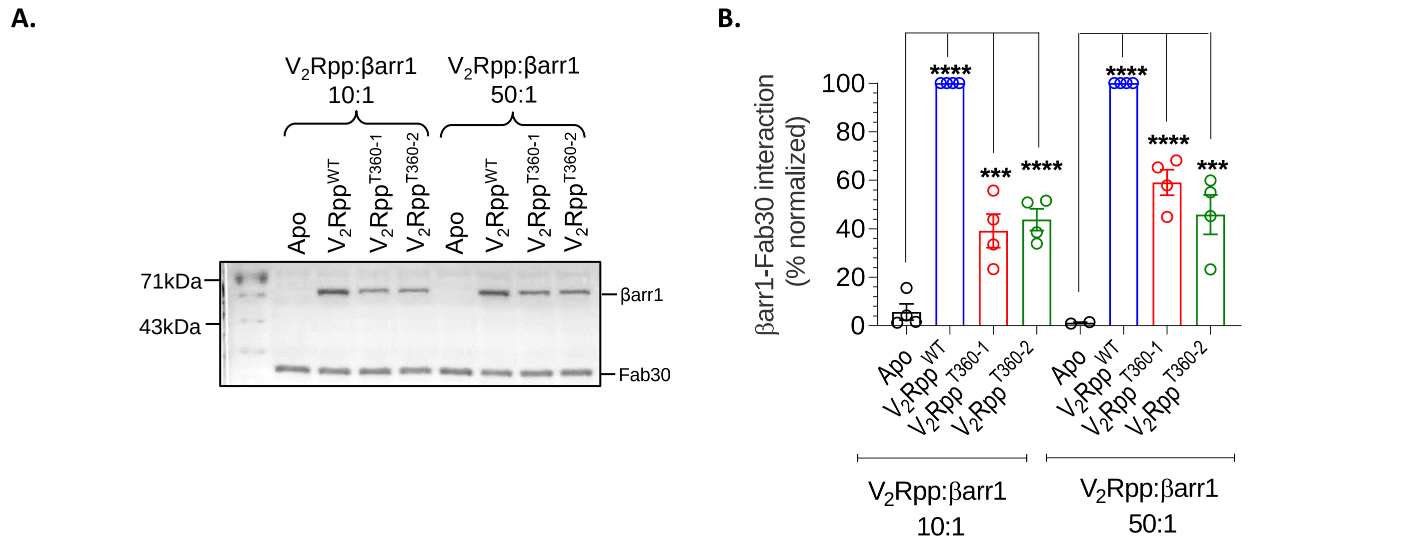


**Figure S2. Recognition of $V_2R^{T360-1}$/ $V_2R^{T360-2}$ activated βarr1 by Fab30 sensor. A.** Co-immunoprecipitation with Fab30 also shows that it recognizes both the $V_2R^{T360-1}$ and $V_2R^{T360-2}$ bound βarr1. $V_2R^{T360-1}$/ $V_2R^{T360-2}$ activated βarr1 was incubated with Fab30, and the complex was then subsequently pulled down with Protein-L agarose beads. A representative blot from four independent experiments is shown here. **B.** Densitometry-based quantification of βarr1- Fab30 interaction normalized with $V_2R^{WT}$-βarr1 control (taken as 100%) and analyzed using one-way ANOVA (*p<0.05, **p<0.01 ***p<0.001, **** p<0.0001) is shown.

**A.**

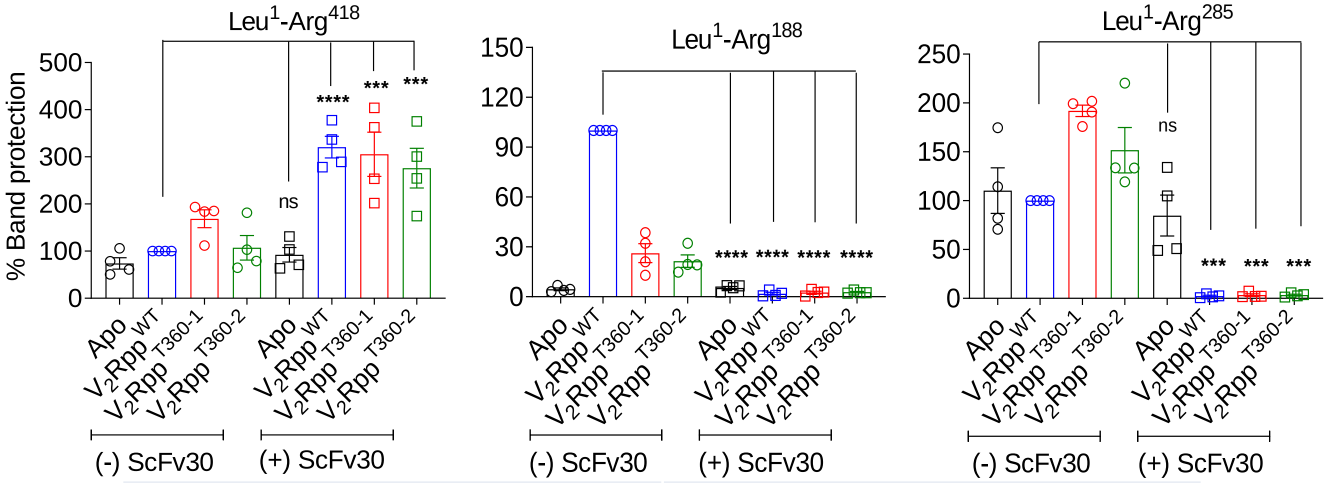


**B.**

| 4JQI | | | 7DFA | | |
|---|---|---|---|---|---|
| Residue | Interacting partner residue | Distance (Å) | Residue | Interacting partner residue | Distance (Å) |
| 188 ARG N | 197 LEU C | 4.00 | 188 ARG CG | 190 A PHE CE1 | 3.68 |
| 188 ARG N | 197 LEU O | 2.90 | 188 ARG CG | 190 A PHE CZ | 3.63 |
| 188 ARG CA | 197 LEU O | 3.84 | 188 ARG NE | 190 A PHE CE1 | 3.97 |
| 188 ARG C | 197 LEU N | 3.90 | 188 ARG NH2 | 341 A SER CA | 3.95 |
| 188 ARG O | 196 PRO CA | 3.42 | 188 ARG NH2 | 340 A SER O | 3.17 |
| 188 ARG O | 196 PRO C | 3.58 | | | |
| 188 ARG O | 197 LEU N | 2.77 | | | |
| 188 ARG O | 197 LEU CA | 3.73 | | | |
| 188 ARG O | 197 LEU O | 3.76 | | | |
| 188 ARG O | 197 LEU CB | 3.81 | | | |
| 188 ARG O | 190 PHE N | 3.99 | | | |
| 188 ARG O | 196 PRO CB | 3.72 | | | |
| 188 ARG CB | 197 LEU O | 3.94 | | | |
| 188 ARG CB | 343 VAL CG2 | 3.57 | | | |
| 188 ARG CG | 190 PHE CZ | 3.57 | | | |
| 188 ARG CG | 343 VAL CG2 | 3.83 | | | |
| 188 ARG CG | 190 PHE CE1 | 3.52 | | | |
| 188 ARG NH1 | 341 SER C | 3.84 | | | |
| 188 ARG NH1 | 341 SER O | 3.84 | | | |
| 188 ARG NH1 | 342 ASP C | 3.89 | | | |
| 188 ARG NH1 | 342 ASP O | 2.78 | | | |
| 188 ARG NH2 | 340 SER O | 3.07 | | | |
| 285 ARG N | 283 A GLU C | 3.43 | 285 ARG CB | 130 GLN O | 3.98 |
| 285 ARG N | 283 A GLU O | 3.13 | 285 ARG CG | 130 GLN O | 3.55 |
| 285 ARG CA | 130 A GLN O | 3.45 | 285 ARG CG | 132 GLY N | 3.66 |
| 285 ARG C | 130 A GLN O | 3.58 | 285 ARG CD | 132 A GLY CA | 3.85 |
| 285 ARG C | 287 A LEU N | 3.27 | 285 ARG CD | 132 A GLY N | 3.55 |
| 285 ARG O | 336 A GLY CA | 3.91 | 285 ARG CD | 135 A ASP OD2 | 3.31 |
| 285 ARG O | 287 A LEU N | 3.55 | 285 ARG NE | 135 A ASP CG | 3.70 |
| 285 ARG O | 335 A LEU C | 3.80 | 285 ARG NE | 135 A ASP OD1 | 3.53 |
| 285 ARG O | 335 A LEU O | 3.47 | 285 ARG NE | 135 A ASP OD2 | 3.04 |
| 285 ARG O | 338 A LEU CG | 3.51 | 285 ARG CZ | 333 A GLY O | 3.58 |
| 285 ARG O | 338 A LEU CD1 | 3.50 | 285 ARG NH1 | 336 A GLY CA | 3.58 |
| 285 ARG CB | 130 A GLN O | 3.89 | 285 ARG NH1 | 336 A GLY N | 3.89 |
| 285 ARG CG | 130 A GLN O | 3.26 | 285 ARG NH1 | 333 A GLY O | 3.47 |
| 285 ARG CG | 131 A PRO C | 3.86 | 285 ARG NH2 | 135 A ASP OD1 | 3.82 |
| 285 ARG CG | 132 A GLY N | 3.61 | 285 ARG NH2 | 333 A GLY O | 3.04 |
| 285 ARG CD | 336 A GLY CA | 3.80 | | | |
| 285 ARG CD | 333 A GLY O | 3.29 | | | |
| 285 ARG CD | 336 A GLY N | 3.88 | | | |
| 285 ARG NE | 333 A GLY O | 3.41 | | | |
| 285 ARG CZ | 333 A GLY O | 3.64 | | | |
| 285 ARG NH1 | 135 A ASP CG | 3.99 | | | |
| 285 ARG NH1 | 135 A ASP OD1 | 3.85 | | | |
| 285 ARG NH1 | 135 A ASP OD2 | 3.33 | | | |
| 285 ARG NH1 | 333 A GLY O | 3.72 | | | |

**Figure S3. A. Trypsin proteolysis pattern of $V_2R^{T360-1}$/ $V_2R^{T360-2}$ activated βarr1 becomes nearly-identical to that of $V_2R^{WT}$ in the presence of ScFv30. A.** Densitometry based quantification of trypsin fragments corresponding to 47kDa ($Leu^1$-$Arg^{418}$), 32kDa ($Leu^1$-$Arg^{285}$) and 21kDa ($Leu^1$-$Arg^{188}$ is shown. Data from four independent experiments, normalized with respect to $V_2R^{WT}$- without ScFv30 condition, and analyzed using one-way ANOVA is presented here. (***p<0.001, **** p<0.0001). **B.** Inter residue contacts (within 4Å) corresponding to residues $Arg^{188}$ and $Arg^{285}$ of 4JQI and 7DFA; calculated separately using CONTACT/ACT program within the CCP4 suite (15).

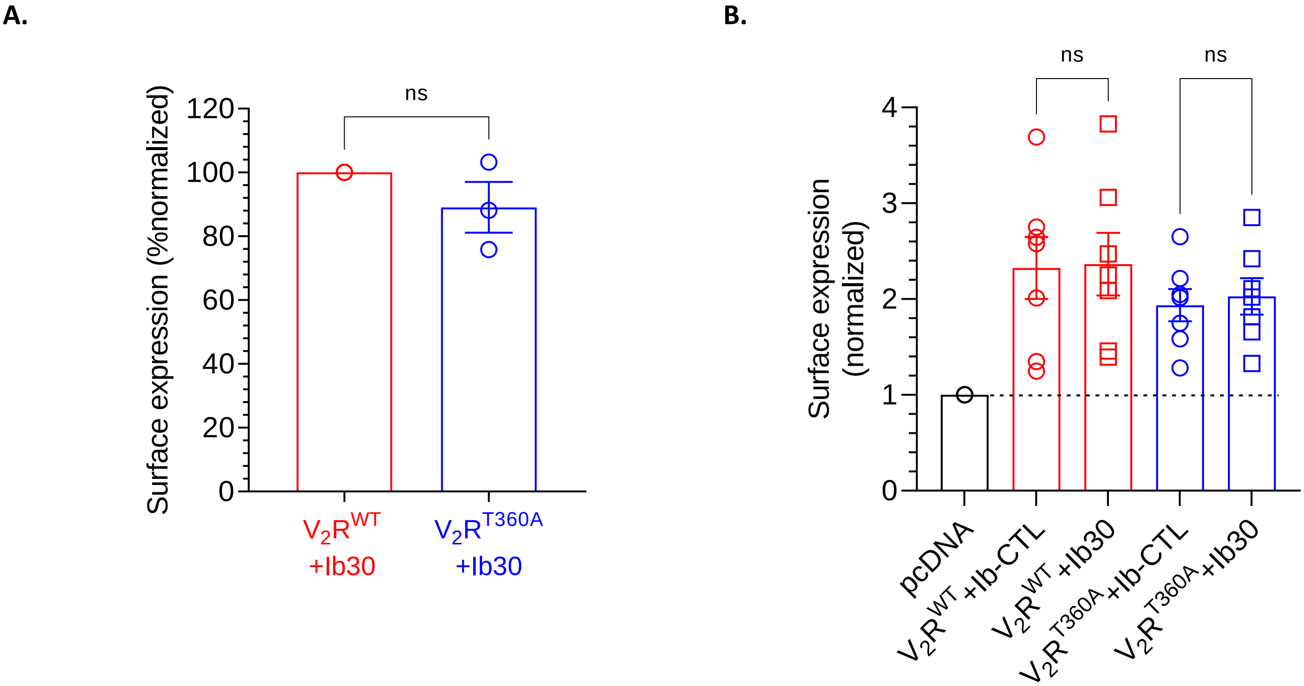


**Figure S4. Surface expression of $V_2R$ constructs in different assays. A.** Surface expression of $V_2R$ constructs in the NanoBiT assay as measured using whole cell ELISA, normalized with respect to $V_2R^{WT}$ (treated as 100%), and analyzed using paired t-test (ns, non-significant). **B.** Surface expression of $V_2R$ constructs in ERK1/2 phosphorylation assays as measured using whole cell ELISA, normalized with respect to vector-transfected cells (treated as 1), and analyzed using one-way ANOVA (ns, non-significant).

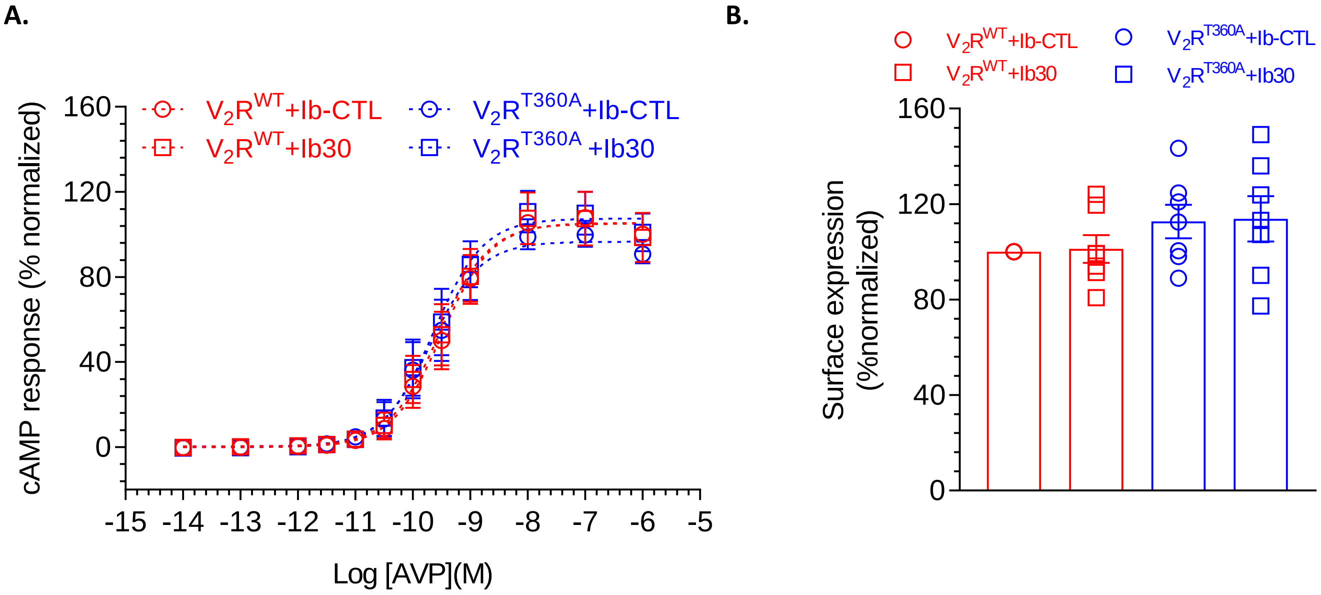


**Figure S5. Effect of Intrabody30 (Ib30) on agonist-induced cAMP response. A**. Ib30 does not affect agonist-induced cAMP response for the $V_2R$ constructs as assessed using GloSensor assay. HEK-293 cells expressing the indicated constructs were stimulated with AVP followed by measurement of cAMP using luminescence-based readout. Data (average±SEM) from seven independent experiments are presented here. **B.** Surface expression of $V_2R$ constructs in GloSensor assay as measured using whole cell ELISA.
